# The effect of spatial bottleneck on human eco-cultural range expansion

**DOI:** 10.1101/2025.10.01.679531

**Authors:** Ryuichi Kumata, Seiji Kadowaki, Joe Yuichiro Wakano

## Abstract

The spatial spread of humans and culture is shaped by feedback between demography and cultural accumulation, as captured by the eco-cultural range expansion model. Yet, the effect of the geographical boundaries on this process remains poorly understood. Here, we extend the model to a two dimensional space where two large habitats are connected by a narrow corridor (spatial bottleneck) with reflecting boundaries. We find that a wave of ecological invasion is unaffected by the bottleneck, whereas a cultural wave can be blocked by sufficiently narrow corridors. The maximum bottleneck width that prevents propagation depends on the domain shape, and it converges to the threshold value when the habitat to be invaded gets larger. Our results align with mathematical results on a bistable reaction–diffusion equation, particularly mean curvature flow, based on which we provide an approximate formula of the threshold bottleneck width. Applied to the Middle– Upper Paleolithic transition, the findings suggest that while the spread of modern humans was robust to spatial bottlenecks, the expansion of advanced Upper Paleolithic culture could be delayed or halted by narrow corridors. Archaeological records point to potential cases where such geographic bottlenecks constrained cultural dispersal.

## 1 Introduction

Recent archaeological and genetic studies suggest that the dispersal of *Homo sapiens* was shaped not only by biological adaptation but also by cultural processes (Groucutt et al., 2015; Fu et al., 2016; Dennell, 2020). A key context is the Middle to Upper Palaeolithic (MP–UP) transition, when the spread of Upper Palaeolithic stone tool technologies coincided with the extinction of Neanderthals, who had coexisted with earlier *Homo sapiens* (Higham et al., 2014; Hublin, 2015). This archaeological overlap indicates that culture may have played an essential role in both the expansion of modern humans and the extinction of archaic populations.

Cultural evolution theory predicts the mutual interaction between population size and cultural level (Henrich, 2004). According to the population size hypothesis, larger populations are more likely to sustain and accumulate culture through social learning, and in turn, accumulated culture provides advanced technologies that support larger population sizes (Henrich, 2004; Kobayashi and Aoki, 2012; Mesoudi and Thornton, 2018). This feedback between population size and cultural level can create bistability between small populations with low level of culture and large populations with high level of culture (Gilpin et al., 2016). Building on this feedback, the eco-cultural range expansion model was proposed to explain how demographic process and cultural expansion interact during human dispersal (Wakano et al., 2018; Wakano and Kadowaki, 2021). The model predicts two types of waves in human expansion. The first, a monostable wave, advances by replacing an unstable equilibrium with a stable one—corresponding to the replacement of archaic populations by a coexistence of archaics and moderns with low cultural level. The second, a bistable wave, replaces one stable equilibrium with another—namely, the replacement of this coexistence by a large modern population with high cultural level. This mechanism may account for archaeological observations: the initial colonization of *Homo sapiens* into Eurasia produced a transient coexistence with Neanderthals, followed by their extinction as *Homo sapiens* populations with high cultural levels expanded during the Middle-to-Upper Paleolithic transition (Wakano et al., 2018; Wakano and Kadowaki, 2021).

The geographical dispersal of human populations was constrained by uninhabitable regions such as seas, mountain ranges, rivers, and deserts (Roberts and Stewart, 2018). As a result, humans often expanded through spatial bottlenecks formed by these geographic barriers. Examples of such bottlenecks include the Sinai penninsula, which connects Africa and Eurasia (Bae et al., 2017). Previous large-scale simulation studies, typically based on complex domains derived from global paleoclimate models, have explored human dispersal under such settings (Timmermann, 2020; Timmermann et al., 2024). However, these studies have commonly employed monostable reaction terms where the effect of spatial bottlenecks is relatively simple.

From a mathematical perspective, boundary conditions and domain shapes critically influence the qualitative behavior of reaction–diffusion systems. This sensitivity is particularly pronounced in systems with bistable reaction terms, where domain shape can fundamentally alter the behaviours of the models. For example, stable non-uniform stationary solutions can emerge in non-convex domains (Matano, 1979; Matano and Mimura, 1983). Thus, since eco-cultural range expansion is essentially a bistable phenomenon, it is crucial to explicitly investigate how geographic bottlenecks shape its dynamics.

Here, we examine how spatial domain affects eco-cultural range expansion. We investigate our eco-cultural range expansion in the specific spatial domains where two large habitats are connected by a narrow habitat with the reflecting boundary condition. We show that narrow bottlenecks can slow the second bistable wave without halting the geographic spread of humans. If the bottleneck is sufficiently narrow, second wave associated with high skilled culture is blocked at its exit. The critical bottleneck width that blocks culture is negatively related to the speed of second wave, consistent with mean curvature flow analysis. We discuss possible archaeological cases where such delays or blockings might have occurred.

## 2 Model

### 2.1 Basic model

To investigate the impact of spatial structure on the spread of human population and their cultures in the geographical space, we extend the previous eco-cultural range expansion model (Wakano et al., 2018; Wakano and Kadowaki, 2021) to that in two dimensional spatial domain Ω:

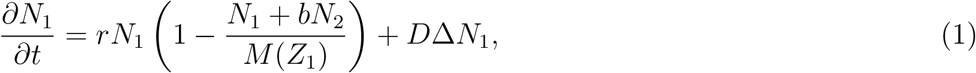

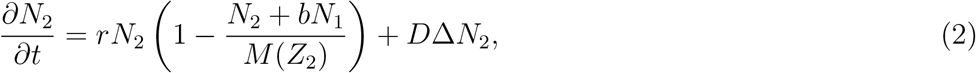

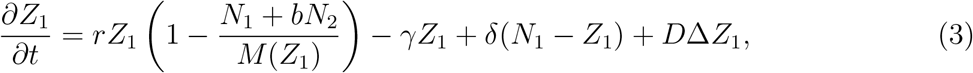

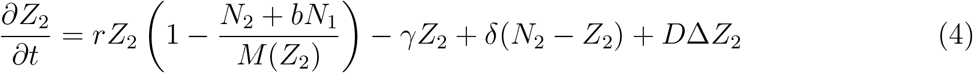

with Laplacian 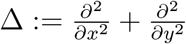 and with zero-flux boundary condition:

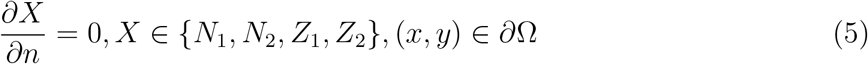

where 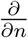 is the derivative in the direction perpendicular to the boundary *∂*Ω (the normal direction), representing the gradient along the normal vector. Our boundary condition assumes that humans do not migrate outside a certain habitable area across the boundary and that humans do not die because of migrating out of the habitable area.

The other variables and parameters follow the same assumptions as in (Wakano et al., 2018). *N*_*i*_ represents the density of species *i* and *Z*_*i*_ represents that of the skilled individuals of species *i. r* is the intrinsic population growth rate of humans and *M* (*Z*_*i*_) is the carrying capacity which depends on the density of skilled individuals

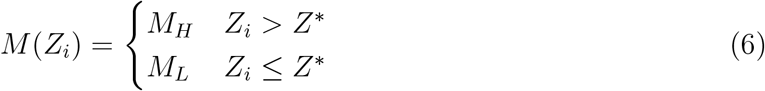

*b* is the interspecific competition coefficient. *D* is the diffusion coefficient of the hominoids. δ is the rate of skill-acquiring and *γ* is the forgetting rate of the skills. Index 1 and 2 refer to the resident archaics and the intrusive moderns. The carrying capacity depends on the density of skilled individuals (population size hypothesis) (Henrich, 2004). Note that each parameter (*r, b, M*_*L*_, *M*_*H*_, *Z*^∗^, δ, *γ, D*) is the same in both species reflecting our assumption that two species has the same cognitive capability and ecological characteristics (i.e., the same reproductive rate, mobility, learning parameters).

### 2.2 Spatial domain

The spatial domain Ω consists of Ω_1_, Ω_2_ and Ω_SB_:

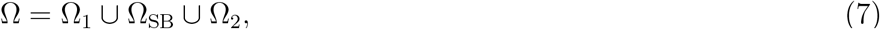

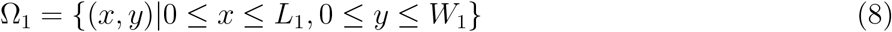

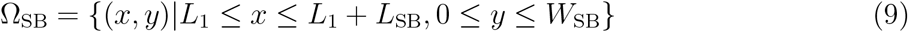

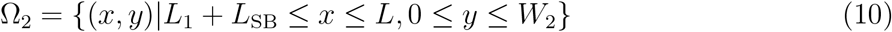

(see Fig. 1). Ω_1_ and Ω_2_ are rectangular regions with lengths *L*_1_ and *L*_2_ in the *x*-direction and widths *W*_1_ and *W*_2_ in the *y*-direction, respectively. They are linked by a narrow corridor Ω_SB_. Ω_SB_ is an elongated rectangular region, representing the spatial bottleneck, with width *W*_SB_ and length *L*_SB_. We define by *P*_1_ and *P*_2_ the points exactly in the middle of the subdomains Ω_1_ and Ω_2_ (Fig. 1). In this study, we introduce the modern humans to a part of Ω_1_ and investigate how they and their culture spread into the areas already occupied by existing archaics.

**Figure 1:**
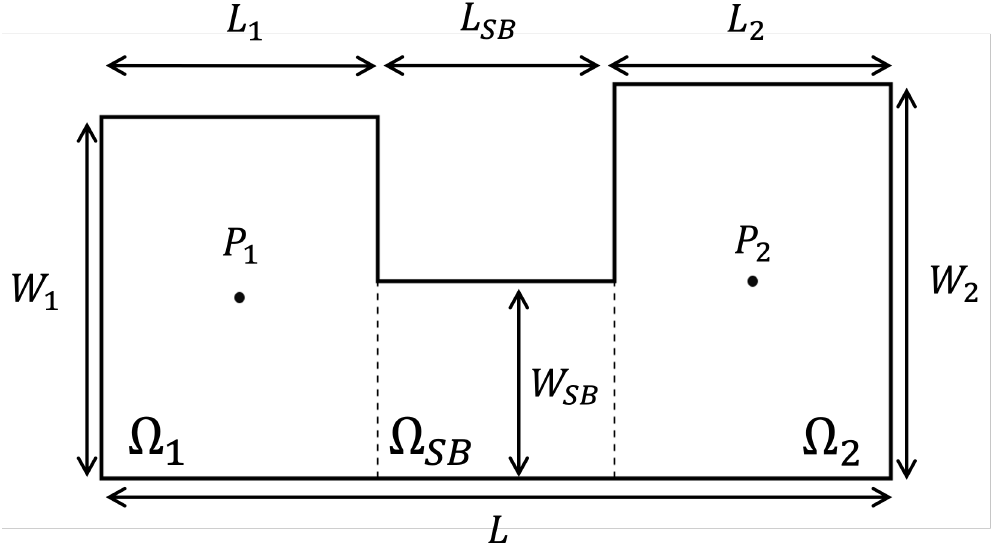
Spatial domain Ω. Two patches Ω_1_ and Ω_2_ are connected by a narrow corridor Ω_*SB*_. *L*_*i*_ and *W*_*i*_ are the length (horizontal) and width (vertical) of Ω_*i*_. *L* = *L*_1_ + *L*_*SB*_ + *L*_2_ is the total length of the domain. *P*_1_ = (*x*_1_, *y*_1_) = (*L*_1_*/*2, *W*_1_*/*2) and *P*_2_ = (*x*_2_, *y*_2_) = (*L* − *L*_2_*/*2, *W*_2_*/*2) indicate the points located at the center of Ω_1_ and Ω_2_.

### 2.3 Scaling time and space

To simplify the analysis, we consider the following rescaling:

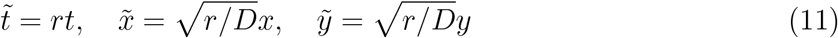

This scale transformation normalizes the diffusion coefficient to 1 and the intrinsic rate of human population growth to 1. With this scale transformation, the above systems are transformed to the following

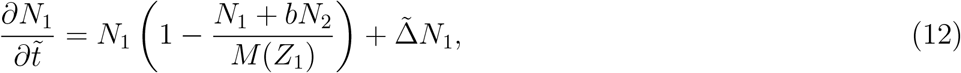

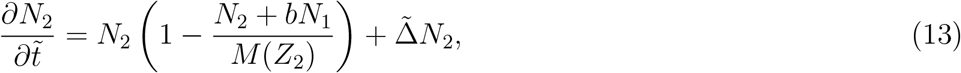

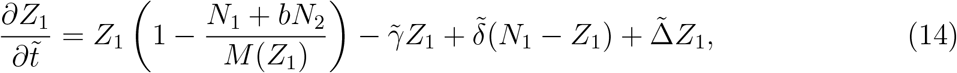

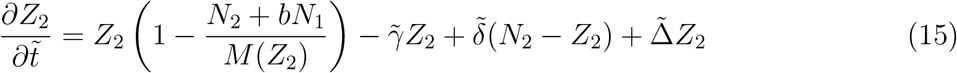

where tilde means the scale transformation and the spatial domain is also scaled to 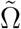. In order to avoid complications in the notation, the tilde symbol is omitted hereafter.

### 2.4 Scaling density

Consider the transformation of the variables

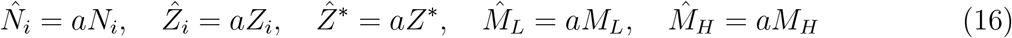

then the equations with and without hat become identical. This means that the model is invariant with the change of the unit of counting population density. To be more specific, the solution of a model with carrying capacity changing from *M*_*L*_=20 [people/km^2^] to *M*_*H*_ =50 [people/km^2^] at critical skilled density *Z*^∗^=15 [people/km^2^] is simply the halved population densities of the solution of a model with carrying capacity changing from *M*_*L*_=40 [people/km^2^] to *M*_*H*_ =100 [people/km^2^] at critical skilled density *Z*^∗^=30 [people/km^2^]. Mathematically, this invariance of the dynamics under the scale transformation of density means that only the ratio *M*_*H*_: *Z*^∗^: *M*_*L*_ is meaningful for the dynamical outcomes of the system.

### 2.5 Reduced system

If the acquisition and loss of cultural skills are faster than the time scale of human reproduction and death, the skilled densities quickly converge to a local cultural equilibrium

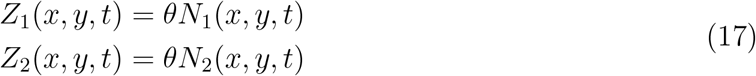

where

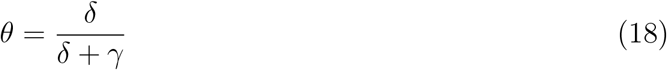

represents the proportion of skilled individual at the cultural equilibrium. Moreover, if (17) is satisfied at *t* = *t*_1_, then it holds for any *t* ≥ *t*_1_ (i.e., it is an invariant manifold of the original four-variable system) (Wakano et al., 2018). In this study we assume this cultural equilibrium and consider the following two-variable system

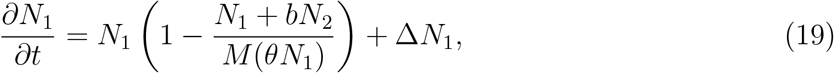

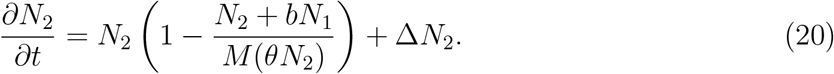

Hereafter, we will study this system.

### 2.6 Equilibria without spatial structure

In the absence of spatial structure, there are in total eight possible equilibrium points with four internal equilibria

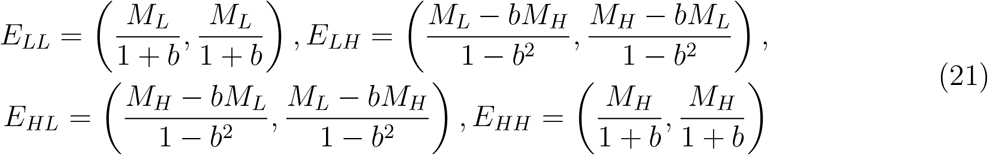

and four edge equilibria

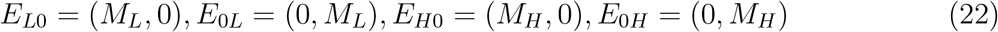

where the first and second quantity in the bracket is the equibrium density of archaic and modern human, respectively. The stability of these equilibrium is classified by the parameter values of the system. If *M*_*L*_ *< Z*^∗^/*θ < M*_*H*_ /(1 + *b*) and 0 *< b < M*_*L*_/*M*_*H*_, the four internal equilibria is stable and the four edge equilibria is unstable. On the other hand, if *M*_*L*_ *< Z*^∗^/*θ < M*_*H*_ /(1 + *b*) and *b > M*_*L*_/*M*_*H*_, the two edge equilibria (*E*_0*H*_, *E*_*H*0_) are stable instead of the two asymmetrical internal equilibria (*E*_*HL*_, *E*_*LH*_). In the latter condition, one human species can be extinct (i.e., at *E*_0*H*_, or at *E*_*H*0_), or the two species can coexist (i.e., at *E*_*LL*_, or at *E*_*HH*_). In the previous study (Wakano et al., 2018), they demonstrated that the model under a one-dimensional spatial structure showed the dynamical replacement of archaic humans by modern humans through the spread of high-density populations of modern humans with the high cultural level (the replacement scenario). Here we only consider the replacement scenario.

### 2.7 Numerical method

#### Numerical simulation of PDE

For numerics, we adapt alternating direction implicit (ADI) method in the finite difference method. We first calculate changes by reaction terms by Δ*t*/2, then calculate the diffusion in the *x*-direction by the implicit method. Next, we do the same for the *y*-direction. We set Δ*x* = Δ*y* = 0.01 (for Fig. 6B) and Δ*x* = Δ*y* = 0.05 (for the other figures). We set Δ*t* = 0.01 (for Fig. 2,Fig. 3AB, Fig. 6B) and Δ*t* = 0.1 (for Fig. 3C, Fig. 5 and Fig. 6A).

**Figure 2:**
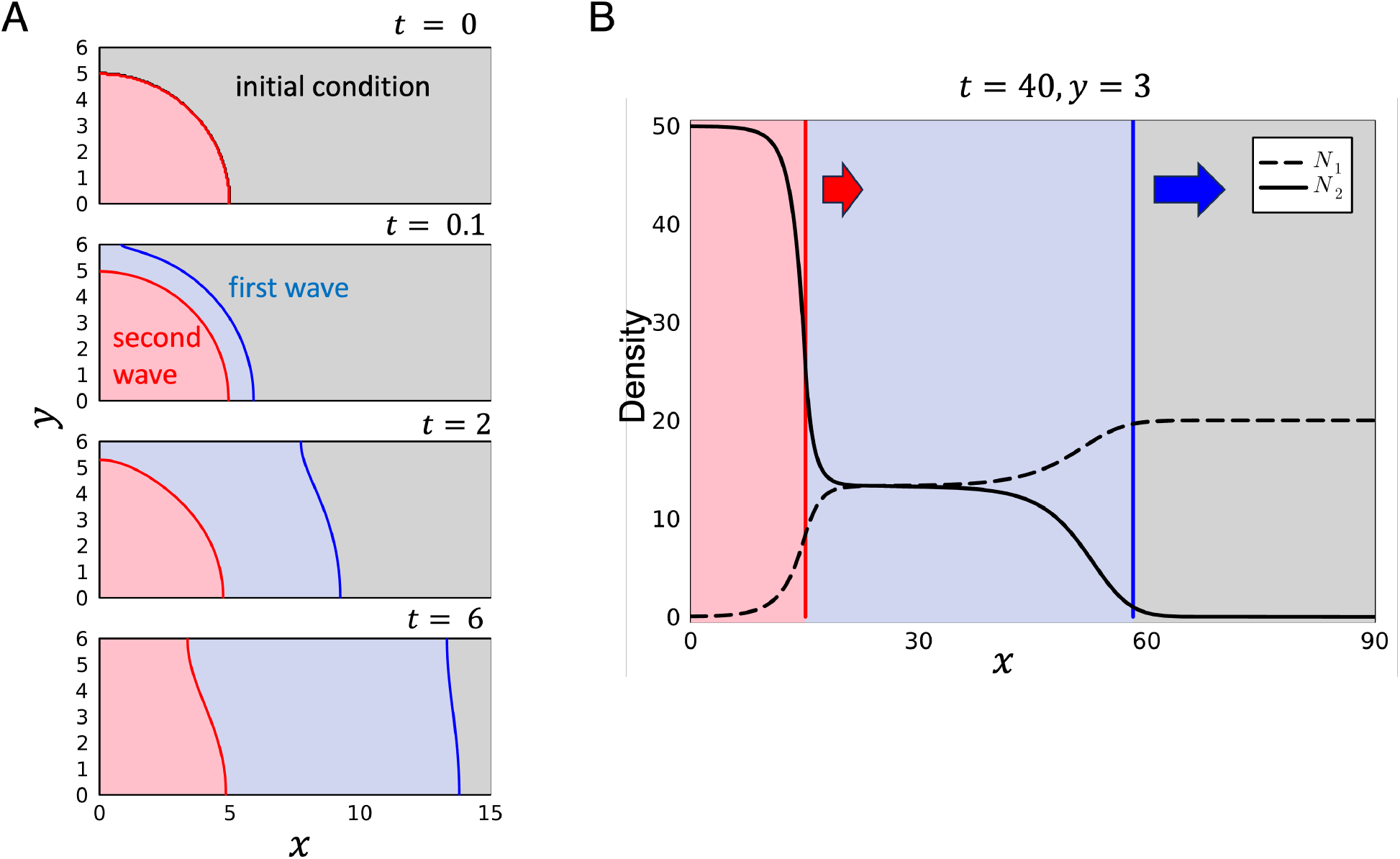
Spatial dynamics of eco-cultural range expansion. A. Snapshots of the distributions. Each panel shows the regions where the modern human density is *N*_2_ *> Z*^∗^/*θ* (light red region), 1 *< N*_2_ *< Z*^∗^/*θ* (light blue region) and *N*_2_ *<* 1 (light gray region) at the indicated time. B. The wave profile at time *t* = 30 and *y* = 3. Black lines show the density of *N*_1_ (dashed) and *N*_2_ (solid). The vertical lines indicate the position of the fronts of the first wave (blue) and the second wave (red). Parameters: *Z*^∗^ = 13, *b* = 0.5, *θ* = 1/2, *M*_*H*_ = 50, *M*_*L*_ = 20. The domain shape: *W*_1_ = 6, *L*_1_ = 90, *L*_SB_ = *L*_2_ = 0

**Figure 3:**
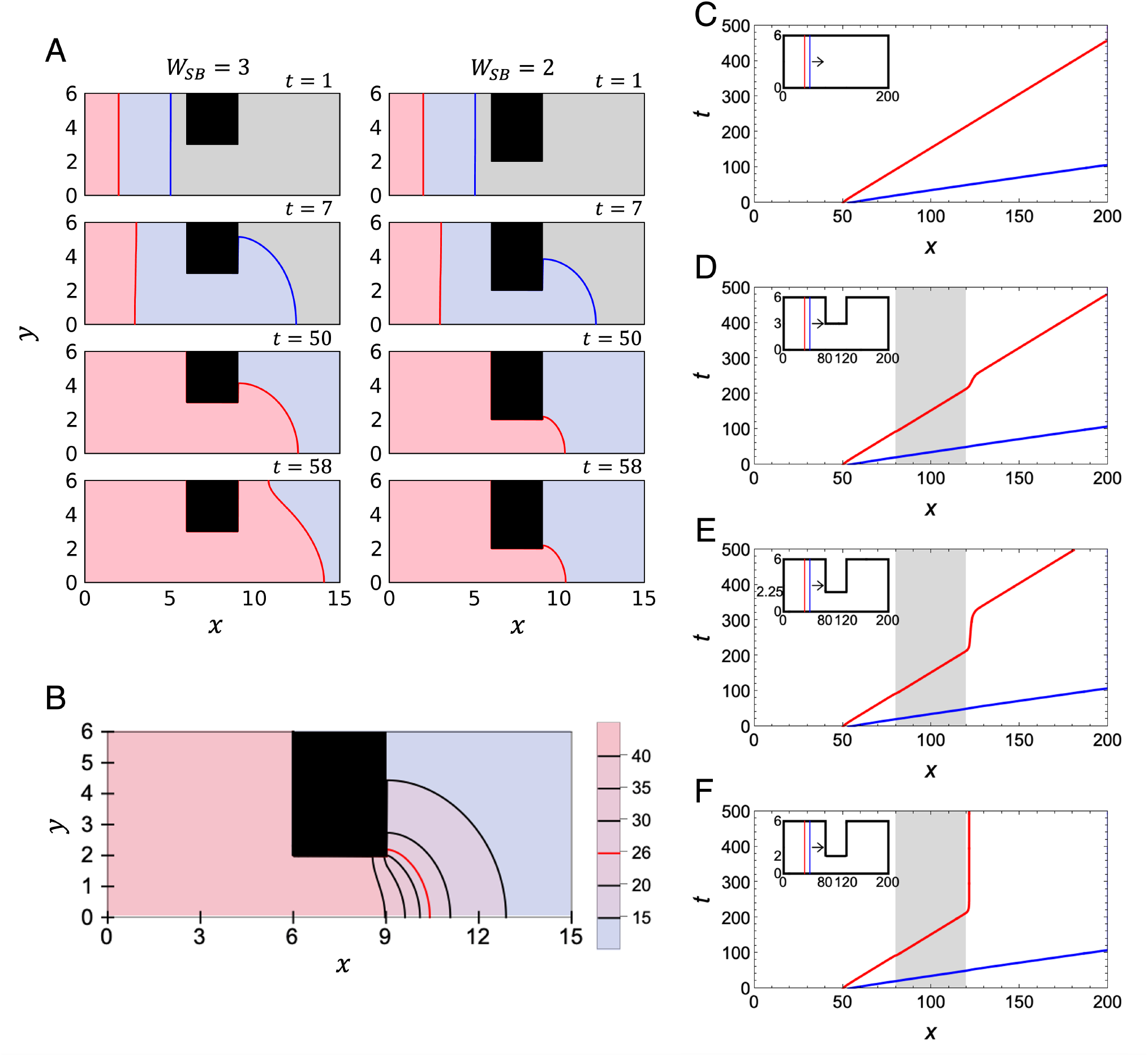
Narrow spatial bottleneck impedes eco-cultural range expansion. Blue curve indicates the position of the first wave (*N*_2_ = 1) and red curve indicate that of the second wave (*N*_2_ = *Z*^∗^/*θ*). A. Snapshots of the distributions at the indicated times with *W*_*SB*_ = 3 (left) and *W*_*SB*_ = 2 (right). B. The density of modern humans *N*_2_ at *t* = 80. Black contours corresponds to *N*_2_ = 40, 35, 30, 20, 15 from left to right. C-F. The position of wave fronts in *x*-axis for the case without bottlenecks (C), with bottleneck width *W*_*SB*_ = 3 (D), with bottleneck width *W*_*SB*_ = 2.25 (E), and with bottleneck width *W*_*SB*_ = 2 (F) in a longer domain. The wave position is calculated at *y* = 1. Gray area represent the region of the bottlenecks. Parameters are the same in the Fig. 2. Domain shape in A and B: *W*_1_ = *W*_2_ = 6, *L*_1_ = *L*_2_ = 6, *L*_SB_ = 3. Domain shape in C: *W*_1_ = *W*_2_ = 6, *L*_1_ = *L*_2_ = 80, *L*_SB_ = 40.

#### Determination of the width of bottleneck to stop the second wave

At every integer time *t* ≥ 1, we calculate 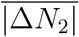, the spatial average of the absolute values of the changes in density of *N*_2_, where Δ*N*_2_(*x, y, t*) = *N*_2_(*x, y, t*) − *N*_2_(*x, y, t*− 1). We stop the simulation at time 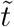 at which it drops below a certain threshold value for the first time 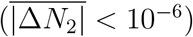. We then determine whether the second wave is blocked or not by comparing the densities of *N*_2_ at the representative points *P*_1_ = (*x*_1_, *y*_1_) and *P*_2_ = (*x*_2_, *y*_2_) at 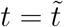 (See Fig. 1). If the densities of *N*_2_ at *P*_2_ is less than 95% of *N*_2_ at *P*_1_:

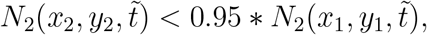

then we judge the second wave is blocked by the bottleneck.

#### Determination of the speed of the second wave

We run the simulation of the system in one-dimensional spatial domain as in (Wakano et al., 2018). We consider the space with lengh 400 with Δ*x* = 0.01 and Δ*t* = 0.1. With the following initial condition

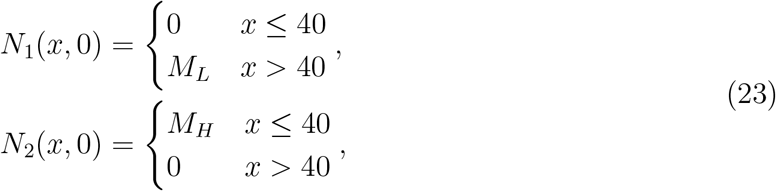

we measured the time and distance from the moment when the front (*N*_2_(*x, t*) = *Z*^∗^/*θ*) of the second wave passed *x* = 200 in the simulation until it exceeded *x* = 280.

## 3 Results

### 3.1 Rectangular spatial domain

Before examining the impact of spatial bottlenecks, we first show the behaviour of our model in the absence of bottlenecks (*W*_1_ = 6, *L*_1_ = 90, *L*_SB_ = *L*_2_ = 0) (Fig. 2) with the following initial conditions:

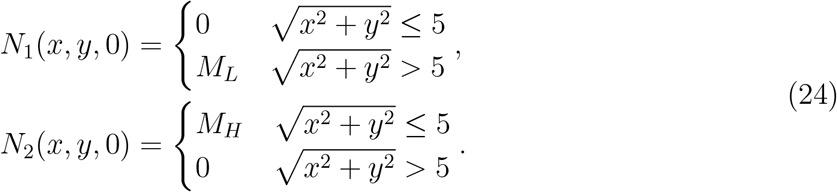

In other words, the modern humans initially only inhabit the circular area with a radius of 5 centered on (0,0) (i.e. the lower left point of Ω) at high density *M*_*H*_, while the rest of the area is occupied only by the archaic humans at low density *M*_*L*_ (Fig. 2A top).

Figure 2A shows the snapshots of the two traveling wave fronts at different times. Similar to the one-dimensional case (Wakano et al., 2018), the modern humans spread out by forming two types of traveling waves. The first wave replaces the unstable equilibrium *E*_*L*0_ (in gray area) where only the archaic humans exist with the stable equilibrium *E*_*LL*_ (in blue area) where the modern and archaic humans coexist at low density. The second wave replaces the stable coexistence equilibrium *E*_*LL*_ (in blue area) with a stable equilibrium where only modern human exists at high density *E*_0*H*_ (in red area). We define the positions of the wavefront according to the density of modern humans. Here, the first wavefront (blue curve) is defined by the position of *N*_2_(*x, y, t*) = 1, and the second wavefront (red curve) is defined by the position of *N*_2_(*x, y, t*) = *Z*^∗^/*θ* at which the carrying capacity switches. These two waves initially spread in two-dimensional space forming spherical waves (*t*=0.1), and when they hit the boundary of *y* = *W*_1_, the shape of the front is distorted and the direction of progression changes. These wave fronts converges to the planar traveling wave solutions that proceed along with x-axis (i.e. the position of the wavefront is no longer dependent on the y-coordinate.).

Figure 2B shows the wave profile (population density of the modern and archaic humans) along with the *x*-axis at *y* = *W*_1_/2 at *t* = 30, which looks identical to that of the one-dimensional case (Wakano et al., 2018).

### 3.2 Effect of spatial bottleneck

We are interested in how the spatial bottlenecks affect eco-cultural range expansion. We consider the areas with longer length in the x-direction than width in the y-direction. In the following, we consider the initial conditions

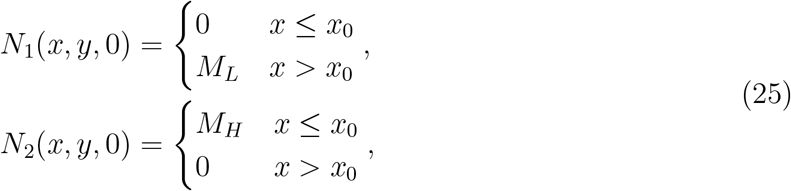

where the population of the moderns with a high culture (*x* ≤ *x*_0_) and the archaic population with a low culture (*x > x*_0_) are separated at the boundary *x* = *x*_0_. Hereafter, we use this initial condition unless we state the other conditions.

Two planer waves were rapidly formed and propagated in the *x*-axis direction in Ω_1_ (*t* = 1 in Fig. 3A). Then they entered the bottleneck Ω_*SB*_ one after another. Figure 3A shows the snapshots of cases with a wider (*W*_*SB*_ = 3) and a narrower (*W*_*SB*_ = 2) bottlenecks. In both cases, the first wave (blue line) went through the bottleneck and propagated in Ω_2_. However, the second wave was blocked at the end of the narrower bottleneck (*W*_*SB*_ = 2). Figure 3B shows the contour plot of the modern human density at *t* = 80 with the narrow bottleneck. Relatively gradual spatial cline was observed at the exit of the bottleneck. To check if this heterogeneous spatial pattern is stationary, we tracked the changes in the distribution of *N*_2_(*x, y, t*) and confirmed that the rate of change was exponentially declining (Fig. 4, see *t >* 80).

**Figure 4:**
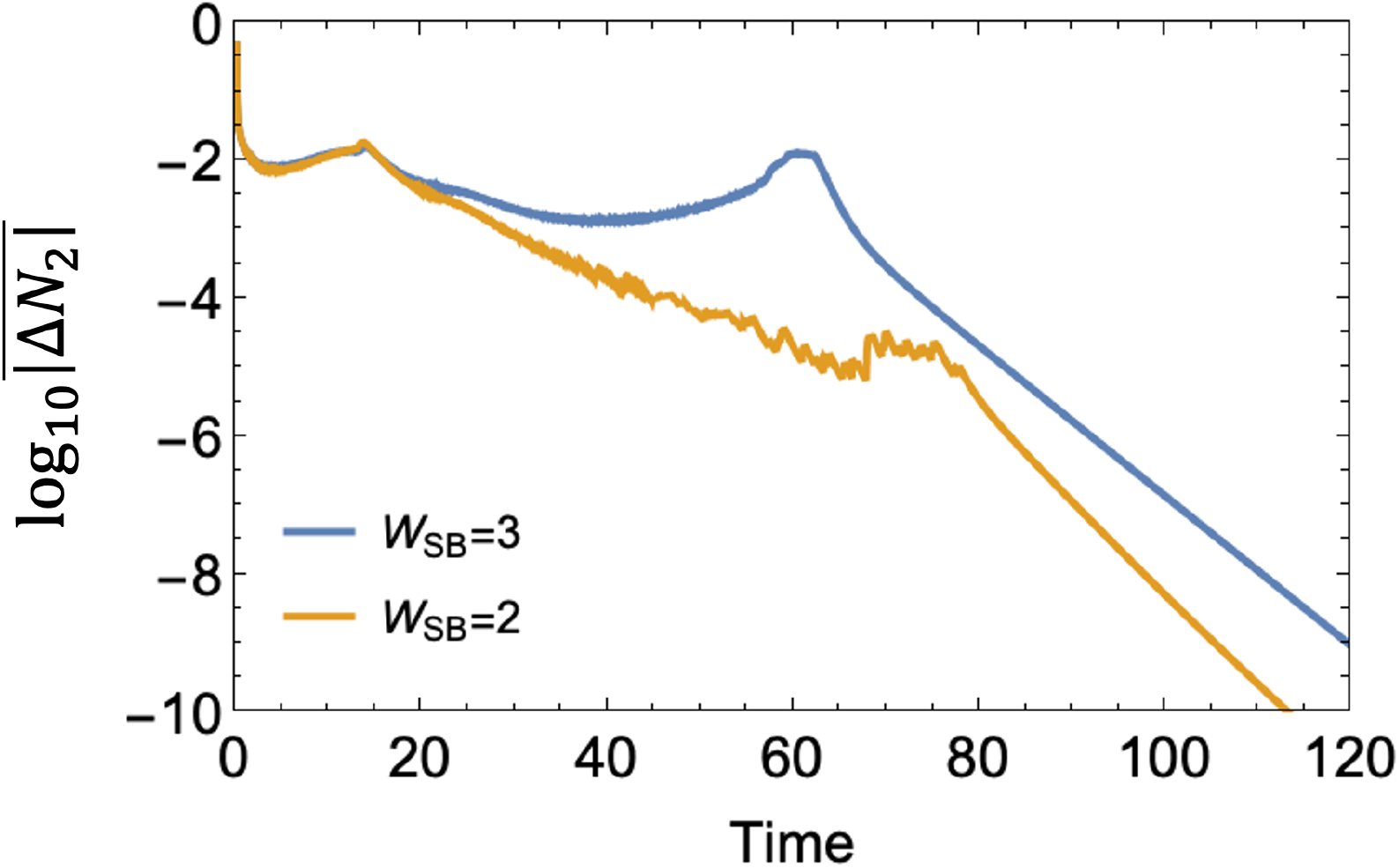
**Dynamical change of the 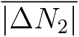 during the eco-cultural range expansion dynamics in Fig. 3A**.

The dynamics of the positions of the two fronts at *y* = 0.75 in four types of large domains (*W*_1_ = 6, *W*_2_ = 6, *L*_1_ = 80, *L*_*SB*_ = 40, *L*_2_ = 80, and *W*_*SB*_ = 2, 2.25, 3, 6) are shown in Fig. 3C-F. The initial condition is given by (25) with *x*_0_ = 50. Without spatial bottleneck (Fig. 3C), the first and second waves quickly reached to their own speeds and the first wave is faster than the second wave. In the case with a wide bottleneck (*W*_*SB*_ = 3), the first wave (blue) was almost unaffected, while the dynamics of the second wave (red) was largely affected by the bottleneck (Fig. 3D). The second wave continued to travel at the same speed inside the bottleneck, but at the exit of the bottleneck it slowed down suddenly at *t* ∼ 200. It took some time until it restarted to travel at *t* ∼ 250. When the wave front departed the bottleneck, the wave recovered the same speed as before and continued spreading in Ω_2_. The delay in progress of second wave at the exit of bottleneck becomes longer with the narrower bottleneck (*W*_*S*_*B* = 2.25, Fig. 3E). For an even narrower bottleneck (*W*_*SB*_ = 2), the second wave did not pass through the bottleneck and came to a complete stop near the end of the bottleneck (Fig. 3F). This is in sharp contrast to the first wave, which is barely affected by the bottleneck.

Numerically observed blocking implies a rapidly-growing waiting time until the second wave restarts or the existence of a stable heterogeneous stationary solution in our system (see Discussion for mathematical details). In the application to human dynamics, they share the same implications: the spatial bottleneck does not block the diffusion of modern humans (first wave) themselves, but it can block the spread of their culture (second wave).

### 3.3 Effect of domain-shape

Our numerics so far suggested the existence the maximum width *ω* such that for any *W*_*SB*_ ≤ *ω* the second wave is blocked. We studied the effect of domain shape on *ω* (Fig. 5). The baseline parameters were *W*_1_ = 24, *W*_2_ = 24, *L*_*SB*_ = 5, *L*_1_ = 12.5, *L*_2_ = 12.5 and the initial values were (25) with *x*_0_ = 5.

**Figure 5:**
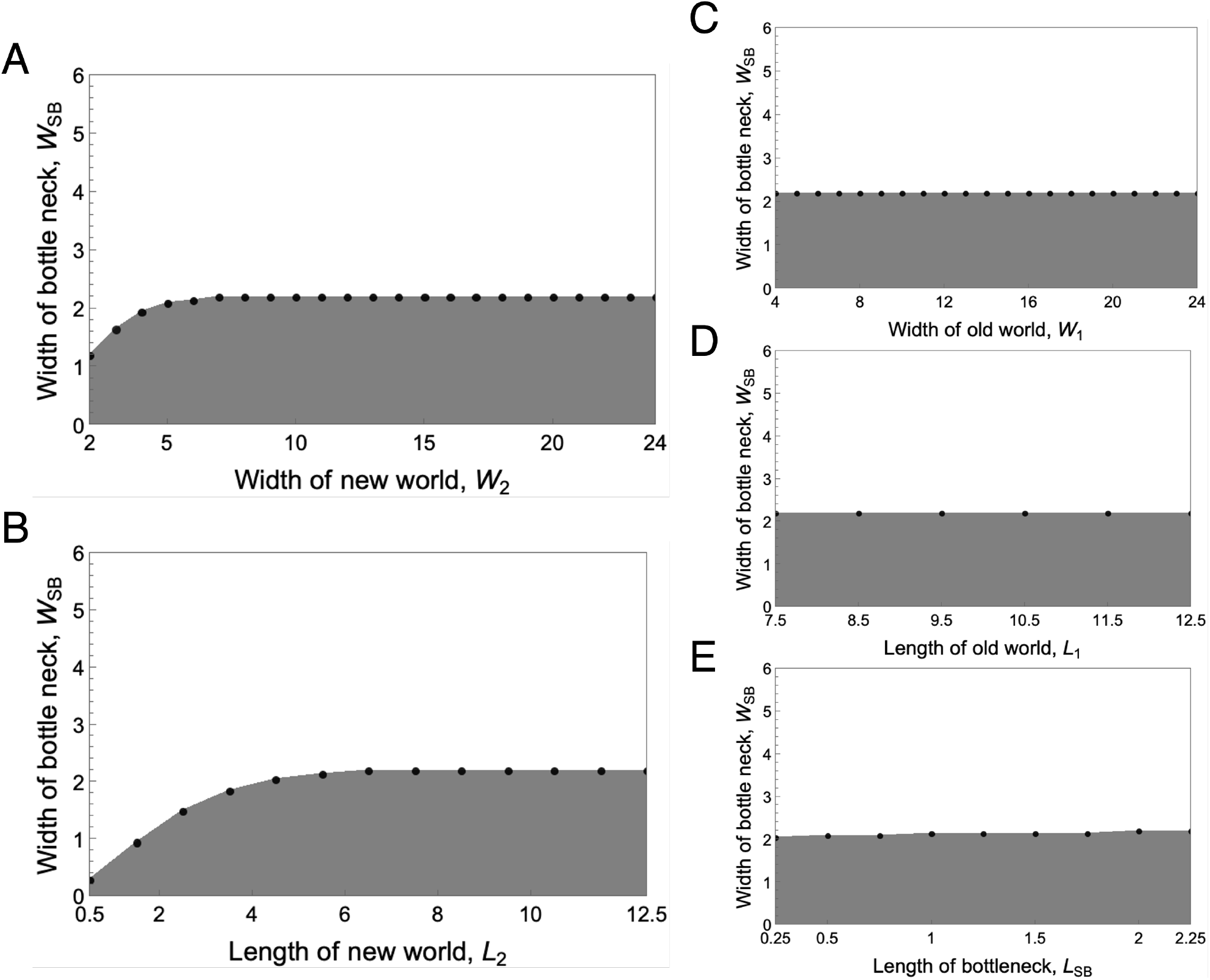
The dependence of domain shape on blocking. Black dot indicates the maximal width *ω* of bottleneck that blocked the second wave. In gray region, the second wave did not pass through the bottleneck. A black diagonal line in A indicates *W*_*SB*_ = *W*_2_, above which Ω_*SB*_ is no longer a bottleneck. Parameters are the same in Fig. 2. The baseline domain shape (if not specified): *W*_1_ = 24, *W*_2_ = 24, *L*_*SB*_ = 5, *L*_1_ = 12.5, *L*_2_ = 12.5

First, we investigated the effect of the domain shape of the new world, Ω_2_ (Fig. 5A). We found that *ω* is an increasing function of *W*_2_ and *L*_2_. Importantly, we found that *ω* converged to the critical width *ω*_*c*_ (around 2.2 under the parameter values used in these simulations) when *W*_2_ and *L*_2_ are sufficiently large. We also found that *ω* hardly depends on the shape of Ω_1_ (i.e., *W*_1_ and *L*_1_), and the length of bottleneck, *L*_*SB*_ (Fig. 5B). In conclusion, we found the critical threshold of bottleneck width *ω*_*c*_ if *W*_2_ and *L*_2_ are sufficient large.

#### The relationship between the speed of second wave and the critical width *ω*_*c*_

In our system, *Z*^∗^/*θ* represents the critical population density that switches the carrying capacity of the population and hence it plays a key role in shaping the speed and direction of the second wave in one-dimensional space. We computed the critical bottleneck width *ω*_*c*_ across a range of *Z*^∗^ values (10.5 ≤ *Z*^∗^ ≤ 14.5 by 0.25) with *θ* = 0.5. As a result, *Z*^∗^/*θ* varied from 21 to 29. We found that *ω*_*c*_ increased with *Z*^∗^/*θ*, indicating that higher *Z*^∗^/*θ* values make the second wave more easily blocked (Fig. 6A). We also measured the speed of the second wave *v*_2_ in the corresponding one-dimensional setting (Fig. 6B). As detailed in the previous study (Wakano et al., 2018), the speed *v*_2_ is a decreasing function of *Z*^∗^/*θ* (Fig. 6C).

**Figure 6:**
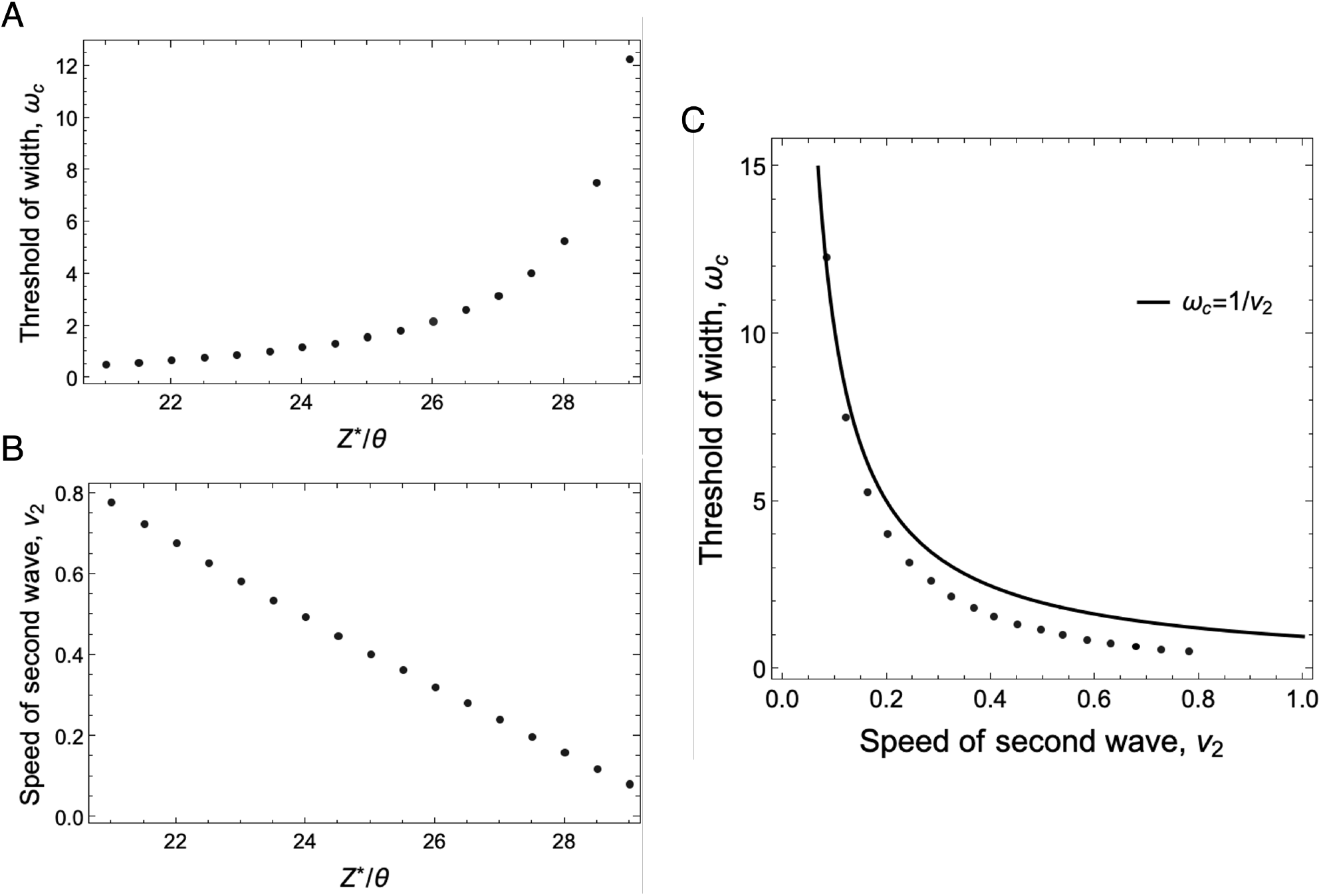
The effect of the speed of the second wave on *ω*_*c*_. A. the dependency of *ω*_*c*_ on *Z*^∗^/*θ*. Black dot indicates the data point of *ω*_*c*_. B. the point is the speed of second wave, *c*, which is numerically determined from the simulation in one-dimensional space. C. The relationship between *ω*_*c*_ and the speed of wave. Black line is given by the mean curvature flow model, *ω*_*c*_ = 1/*v*_2_. The points are obtained by the combination of *ω*_*c*_ and *c* for different *z*^∗^/*θ*. The basic domain shape: *W*_1_ = 24, *W*_2_ = 24, *L*_*SB*_ = 12, *L*_1_ = 24, *L*_2_ = 24. Parameters are the same in Fig. 2 (if not specified).

#### 3.4.1 Mean curvature flow

Previous mathematical studies on traveling waves in reaction-diffusion models give some clues to understand these behaviors. Chen (1992) showed that a traveling front of a bistable one-species system in multi-dimensional space can be described by a mean curvature flow (MCF) model when reaction occurs very quickly so that wave profile becomes very sharp. MCF model when diffusion coefficient is rescaled to *D* = 1 is given by

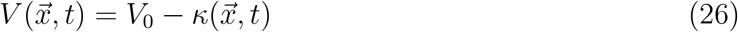

where 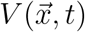 is the normal speed of wave front at position 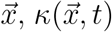 is mean curvature of wave front, and *V*_0_ is a constant determined by a reaction term. By definition, *V*_0_ is the speed of traveling wave in the corresponding one-dimensional system.

Consider a wave front in Fig. 3A. Initially the front is planar so 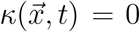. Recall that curvature is the inverse of the radius and the planar front has the infinite radius, i.e., zero curvature. Thus, the normal speed *V* of the second wave is identical to *v*_2_. When the wave is about to exit the bottleneck (see Fig. 3A *t*=50), the wave front will take a round shape whose radius should be around *W*_*SB*_. By assuming a MCF model with the constant curvature *κ* = 1/*W*_*SB*_, we obtain the condition for the second wave to be blocked as

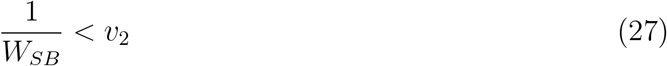

The MCF model predicts a negative relationship between *ω*_*c*_ and *v*_2_ (a curve in Fig. 6C). This prediction agrees well with our numerical results when *ω*_*c*_ is large, but shows noticeable discrepancies when *ω*_*c*_ is small. In cases where MCF approximation works well (*Z*^∗^/*θ* = 28.5, *W*_*SB*_ = 7.6 ∼ *ω*_*c*_) (see Fig. 6A,C), the stationary distribution of *N*_2_ appeared to have a thin circular transitional layer (Fig. 7). In contrast, when the MCF approximation breaks down (*Z*^∗^/*θ* = 26, *W*_*SB*_ = 2 ∼ *ω*_*c*_), the transitional layer appeared thick and distorted rather than being perfectly circular arc (Fig. 3B). This breakdown occurs because the thickness of transitional layer (i.e., the width of wave profile) becomes large relative to *W*_*SB*_, making it impossible to approximate the full dynamics by the dynamics of the wavefront as a thin curve—a key assumption of the MCF model. It is, however, important to note that the thicknesses of the transitional layers for the two cases are similar. They are different relative to the size of spatial domain (compare scales of Fig. 3C and 7). When the spatial domain is large enough (i.e., large *W*_*SB*_), we conjecture that the MCF approximation is good. When *Z*^∗^/*θ* is small and hence *v*_2_ is large, it predicts small *ω*_*c*_ value, which is not large enough for the MCF approximation. This explains why the approximation is not good for small *Z*^∗^/*θ*.

**Figure 7:**
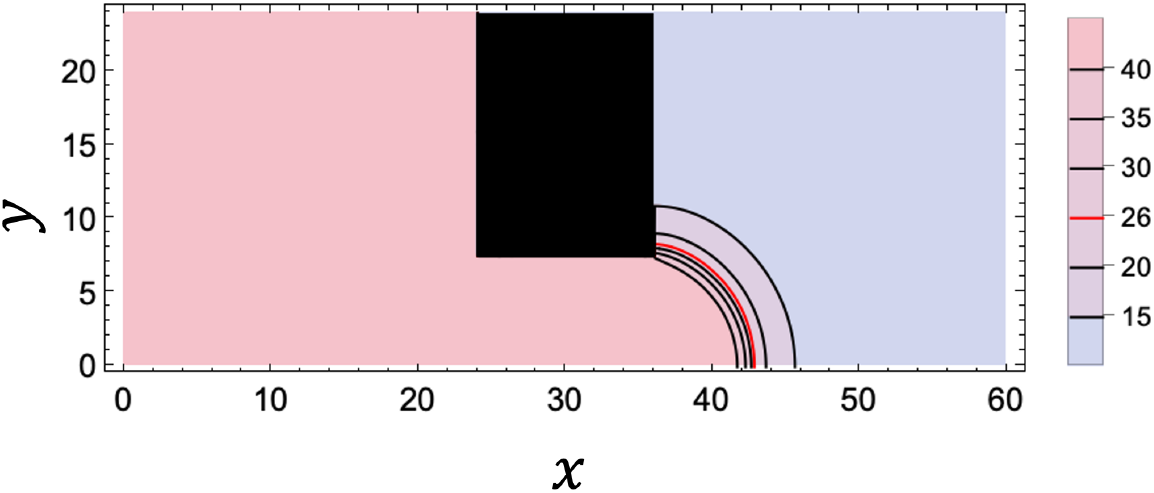
Stable distribution when MCF model gives a good approximation for *ω*_*c*_. The distribution of *N*_2_ when *Z*^∗^/*θ* = 28.5 and *W*_*SB*_ = 7.5 is plotted from the simulation at *t* = 1040, at which the stable condition satisfies 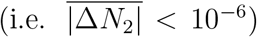. The domain shape is *W*_1_ = 24, *W*_2_ = 24, *L*_*SB*_ = 12, *L*_1_ = 24, *L*_2_ = 24. Parameters are the same in Fig. 2 (if not specified).

### 3.5 Speeds and width with units

We have so far shown results after rescaling time and space so that *r* = *D* = 1. Here we provide a numerical example when we assume parameter values in [km] and [year]. The first wave is of a Fisher-KPP type and the (minimum) speed is given by (Wakano et al., 2018) as

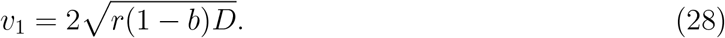

This indicates the speed of geographical spread of modern humans. With our baseline parameter values (*b* = 0.5), we have a unit-less speed of 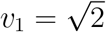. Assuming that one generation of humans is 15 [years] and that the population will double every generation under the ideal condition, we *D*/*r* ≈ 44 [km]. have the intrinsic growth rate *r* = (log_10_ 2)/15 ≈ 0.02 [/year]. Assuming that the inter-specific competition is the half of intra-specific competition (*b* = 0.5) and that a diffusion coefficient is *D* = 10 [km^2^/year], we have the speed of the first wave 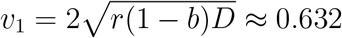 [km/year].

The dynamics of the second wave depends also on the ratio *M*_*L*_: *Z*^∗^/*θ*: *M*_*H*_. Using the parameter values shown in Table 1, a unit-less speed of the second wave *v*_2_ ≈ 0.3 is translated into 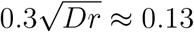 [km/year] and the critical width *ω*_*c*_ ≈ 2 is translated to 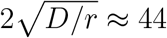[km]. When we increase *Z*^∗^ to 14 [/km^2^], we have the speed ≈0.08 [km/year] and the critical width ≈121 [km]. Recall that these values do not depend on the unit of counting population.

**Table 1:** List of parameters and their units. Values are only for the purpose of explanation.

| Parameter | Value | Unit |
| --- | --- | --- |
| $r$ | 0.02 | /year |
| $\theta$ | 0.5 | — |
| $b$ | 0.5 | — |
| $Z^*$ | 13 | /km<sup>2</sup> |
| $M_L$ | 20 | /km<sup>2</sup> |
| $M_H$ | 50 | /km<sup>2</sup> |
| $D$ | 10 | km<sup>2</sup>/year |

## 4 Discussion

The prehistoric human spread was shaped by a complex process involving the geographical spreads of humans and their culture. In this study, we investigated the impact of spatial bottlenecks on geographical spread of humans and their culture. A key finding is that spatial bottlenecks do not affect the geographical spread of the modern humans (the first wave), but they do hinder the spread of the high-density-high-culture equilibrium and the resulting extinction of archaic humans (the second wave). Spatial bottlenecks slow down or completely block the second wave if they are sufficiently narrow. We characterized the critical width of the bottleneck below which blocking occurs.

### 4.1 Why blocking occurs

The mechanism of blocking can be intuitively understood by the effect of the dilution of population density. Consider the second wave with a planar front with a constant speed *v*_2_ *>* 0 (see the red line in Fig. 3A *t* = 7). It has a monotonic wave profile as shown in Fig. 2B. At each location, modern human density is increasing over time. Since the second wave is a bistable wave, the reaction term is negative in front of the wave and positive behind it. The diffusion term can be negative or positive depending on whether the wave profile is convex or concave in the direction of the travel. Mathematically speaking, in a planar wave we have *∂*^2^*N*_2_/*∂y*^2^ = 0 and thus Δ*N*_2_ = *∂*^2^*N*_2_/*∂x*^2^. The speed *v*_2_ is determined by the balance between the reaction and the diffusion in the *x*-direction. This traveling wave has a positive speed because the positive effect from the diffusion (i.e., migration from high density area behind) overcomes the negative effect from the reaction (i.e., tendency to fall back to the lower equilibrium density) at all locations in front of the wave.

Now consider a system with a spatial bottleneck. When the wave is about to exit the bottleneck, the dilution occurs since human population also diffuses in the *y*-direction. This brings an additional effect from the diffusion term that tends to decrease population density. Mathematically speaking, *∂*^2^*N*_2_/*∂y*^2^ starts to be negative. This makes the second wave tentatively slow. When all the effects are balanced in all spatial points, there appears a spatially heterogeneous stationary solution, and the second wave is blocked (Fig. 3B and Fig. 7).

### 4.2 Implication for Mathematics

If the second wave is blocked at the bottleneck while the first wave propagates to the (right) end of the finite spatial domain Ω, it means that the system has a stable heterogeneous stationary solution. Matano (1979) mathematically proved the existence of a finite domain Ω in which a stable heterogeneous stationary solution exists in a bistable one-species reaction-diffusion equation with zero-flux boundary conditions. Matano and Mimura (1983) obtained the same result for a bistable two-species reaction-diffusion system. For such a solution to exist, Ω needs to be convex, which is consistent with our model (Fig. 1).

In the more recent study, Berestycki et al. (2016) proved that blocking of bistable waves occurs when they go through an abrupt opening (Berestycki et al., 2016). Furthermore, Etheridge et al. (2022) studied an Allen-Cahn equation, a reaction diffusion equation with bistable reaction term, and showed the blocking of the bistable wave occurs by the wide opening domain, which is similar to our domain shape (Etheridge et al., 2022). All these rigorous results are shown for one-species bistable dynamics and our results provide an example of such blocking dynamics in a two-species reaction-diffusion system.

### 4.3 A remark for boundaries

Geographical barriers such as mountains and oceans can be modeled in different ways. Here we assumed the zero-flux boundary condition, where human migration is reflected by the boundaries. Such dynamics appear when human individual or group knows the existence of the barrier and does not try to migrate beyond it. We assumed that habitats are homogeneous and equally habitable inside the boundary. Yet another way to model geographical barriers is to assume less habitable environment for regions like mountains and deserts. Random migration modeled by simple diffusion terms in heterogeneous environment implies that human individual or group keeps migrating into bad environments and dying there. In the real world, there are the effects of spatial heterogeneity and reflecting boundaries. Here we only studied the latter, and the combined effect is a future work.

### 4.4 Implication for archaeology

The results of this study on eco-cultural spatial dynamics offer new insights into the spatial distribution patterns of archaeological cultures, particularly in regions characterized by geographic bottlenecks. One such region is the area around the Sinai Peninsula, which serves as a land bridge connecting the African and Eurasian continents. This corridor is well known as a major route for repeated human dispersals out of Africa since approximately two million years ago. In the case of Lower Paleolithic dispersals, the Oldowan and Acheulean lithic assemblages found in the Levant have been interpreted as evidence of lithic technologies that spread from Africa alongside archaic hominin groups (Bar-Yosef and Belmaker, 2017; Goren-Inbar et al., 1994; Le Tensorer et al., 2015).

Regarding the dispersals of anatomically modern humans (AHMs), the Nubian Levallois technology has been suggested to represent the range expansion of AMHs from Africa to south-west Asia during the middle phase of the Middle Paleolithic (Goder-Goldberger et al., 2016; Oron et al., 2025; Rose et al., 2011). Lithic assemblages characterized by this technology are found across northeastern Africa, the southern Levant, and the Arabian Peninsula. In addition, archaic human populations may have coexisted with AMHs in the Levant during this phase (Hershkovitz et al., 2021; Zaidner et al., 2025), potentially corresponding to the first wave in our eco-cultural range expansion model.

In contrast, the distribution of Early Upper Paleolithic bladelet technology shows a sharp boundary between northeastern Africa and the Levant. This technology, known as the Ahmarian industry, developed in the Levant by AMHs and subsequently spread into the Sinai Peninsula (Bar-Yosef and Belfer, 1977; Marks, 1981; Phillips, 1988). However, it did not extend into northeastern Africa, where other lithic technologies existed that resembled earlier ones in the Levant, such as blade technology Olszewski (2009). This distributional boundary stands in contrast to the widespread presence of similar bladelet technologies across other regions of West Asia, such as the Zagros and the Caucasus, as well as in Europe, where they are represented by the Aurignacian industries (Adler et al., 2008; Ghasidian et al., 2019; Hublin, 2015).

As suggested in our previous studies, bladelet technology in the Early Upper Paleolithic can be regarded as a skill that became widespread during a phase of high population density, as predicted by our eco-cultural range expansion model (Wakano et al., 2018). The halt in its spread at the Sinai Peninsula appears to support the model’s prediction that the cultural phase of high population density tends to terminate at the end of a spatial bottleneck.

Another region potentially relevant to our eco-cultural model is the Iberian Peninsula, where the presence of Protoaurignacian or Early Aurignacian bladelet assemblages is sparse and remains debated, particularly in areas south of the Ebro River basin (Zilhão, 2021). The Protoaurignacian and Early Aurignacian are lithic industries attributed to AMHs and are characterized by bladelet technology (Falcucci et al., 2017). Sites associated with these industries are abundant in the Franco-Cantabrian region, with their southern distribution bounded by the Cantabro-Pyrenean cordillera (Djakovic et al., 2022). The delayed spread of Aurignacian bladelet technology into the Iberian Peninsula has been linked to the prolonged presence of Neanderthals using Middle Paleolithic technology, as proposed in the Ebro Frontier model (Zilhão, 2000). However, the validity of this model remains contested, particularly regarding the chronological evidence for the appearance of Aurignacian assemblages and the persistence of Middle Paleolithic sites (Cortés-Sánchez et al., 2019; Haws et al., 2020; Zilhão, 2021).

Here, we do not intend to engage in the chronological debates surrounding the Ebro Frontier model, but rather focus on its proposition that the delayed appearance of Aurignacian bladelet technology was due to physical barriers such as the Cantabro-Pyrenean cordillera and the Iberian Systems, which impeded the terrestrial movement of Paleolithic foragers (Zilhão, 2021). This idea is broadly relevant to the findings of our study, as our model suggests that spatial bottlenecks formed by mountainous terrains can constrain the spread of the high-population-density cultural phases, such as those represented by the Protoaurignacian and Early Aurignacian.

### 4.5 Further perspective

In this study, we examined a scenario of the Middle-to-Upper Palaeolithic cultural transition. Nevertheless, our eco-cultural range-expansion model can also be applied to other major transitions in which demographic and cultural processes interact. A prominent example is the Neolithic revolution—the transformation from hunting and gathering to farming and herding. Previous models have distinguished between demic diffusion (the spread of farming through migrating populations) and cultural diffusion (the adoption of farming by resident hunter–gatherers), typically formalizing these processes as monostable reaction–diffusion fronts (Fisher-type fronts) (Fort, 2012; 2015). However, the Neolithic transition may also exhibit characteristics indicative of bistable dynamics: farming could be sustained only above a critical population or knowledge threshold. Such threshold effects are readily accommodated within our eco-cultural framework. Extending this approach to the Neolithic case may thus provide novel insights, for example into the ways spatial bottlenecks constrained—or facilitated—the spread of early farming societies.

## Acknowledgements

All authors are supported by MEXT KAKENHI No.24H00001. We thank Kota Ikeda and Ryunosuke Mori for discussion about mathematics.

